# γδ-Thymocyte Maturation and Emigration

**DOI:** 10.1101/2021.03.19.435877

**Authors:** K. Joannou, D.P. Golec, H. Wang, L.M. Henao-Caviedes, J.F. May, R.G. Kelly, R. Chan, S.C. Jameson, T.A. Baldwin

## Abstract

The thymus is the site of both αβ and γδ-T cell development. After several unique waves of γδ-T cells are generated in, and exported from, the fetal/neonatal thymus, the adult thymus continues to produce a stream of γδ-T cells throughout life. One intriguing feature of γδ T cell development is the coordination of differentiation and acquisition of effector function within the fetal thymus, however, it is less clear whether this paradigm holds true in adult animals. To investigate the relationship between maturation and time since V(D)J recombination in adult-derived γδ-thymocytes, we used the Rag2pGFP model. Immature (CD24^+^) γδ-thymocytes expressed high levels of GFP while only a small minority of mature (CD24^-^) γδ-thymocytes were GFP^+^. Similarly, most GFP^+^ γδ-splenocytes were immature, while some were mature. Analysis of γδ-recent thymic emigrants (RTEs) indicated that most γδ-T cell RTEs were CD24^+^ and GFP^+^ and adoptive transfer experiments showed that immature γδ-thymocytes could be maintained in the periphery for at least 3 days over which time they matured. With respect to the mature γδ-thymocytes that were GFP^-^, parabiosis experiments demonstrated that mature γδ-T cells did not recirculate from the periphery. Instead, a population of mature γδ-thymocytes remained resident in the thymus for at least 60 days while mature γδ-thymocytes derived solely from adult hematopoiesis were mostly lost from the thymus within 60 days. Collectively, these data demonstrate two streams of actively developing γδ-T cells in adult mice: an immature subset that quickly leaves the thymus and matures in the periphery, and one that completes maturation within the thymus over a longer period of time. Furthermore, there is a fetal-derived and heterogeneous population of resident γδ-thymocytes of unknown functional importance.

## Introduction

Although γδ-T cells constitute a relatively minor fraction of total T cells, their numerous activities in diverse tissues play important roles in organism-wide immunity. γδ-T cells are present at especially low frequencies in blood, and secondary lymphoid tissue, but are particularly enriched in barrier tissue sites, such as in the gut, lung, uterus, tongue and skin. For reasons that are not yet clear, the development of these tissue specific γδ-T cells occurs in distinct waves during fetal development, with the expression of particular Vγ chains being associated with waves that correspond to homing to specific tissues ^**1**,**2**^, and effector functions as IL-17A or IFNγ producers^**3**^. Tonegawa nomenclature is used for Vγ segments in this article^**4**^. Only Vγ1.1 and Vγ2 expressing and IFNγ producing or naive-like γδ-T cells continue to be produced throughout adulthood in mice^**5**^. These adult-derived γδ-T cells form a circulating population, whereas waves of fetal-derived γδ-T cells quickly become resident in tissues after maturation^**6**^.

The thymus provides the environmental context needed to induce thymocyte progenitors to produce a T Cell Receptor (TCR) through V(D)J recombination. Expression of the γδ-TCR first occurs in the most immature, CD24^+^, progenitors. Similar to its use in tracking αβ-T cell development, downregulation of CD24 correlates with the capability of γδ-thymocytes to produce cytokines^**7**^. In addition to CD24, the expression of CD73 is a useful marker for staging γδ-thymocyte development. In the fetal stage, induction of CD73 occurs following down-regulation of CD24 in IL-17 producing γδ-thymocytes^**8**^, while in adults, expression of CD73 marks commitment to the γδ lineage, and is associated with TCR signaling^**9**^. Thus, it is useful to define γδ-thymocytes in adults as immature CD73^-^ (CD24^+^CD73^-^), immature CD73^+^ (CD24^+^CD73^+^), or mature (CD24^-^CD73^+^). Similar to conventional αβ-T cells, expression of the transcription factor KLF2 and homing receptor S1P1 have been shown to regulate γδ-thymocyte emigration^**10**^. Although the general sequence of γδ-T cell maturation has been elucidated, the coordination of maturation and thymic emigration remains poorly understood. The cellular context of γδ-T cells at different stages of development is a critical question, as the signals a cell receives from its environment direct lineage specification and maturation.

A particularly intriguing feature of γδ-thymocyte development is the apparent coordination of lineage specification and acquisition of effector function while in the thymus. In contrast to conventional αβ-T cells that exit the thymus in a naïve, undifferentiated state, γδ-thymocytes have been shown to express key transcription factors that specify lineage^**11**^, and are capable of rapid cytokine production given stimulation^**5**^. Collectively, this has led to the notion that these cells exit the thymus in a mature state. However, most of these studies have focused on fetal γδ-thymocytes^**5**,**8**^.

Contradicting the notion that only mature γδ-thymocytes emigrate from the thymus, studies performed in adult mice suggest that relatively immature progenitors that lack the ability to produce effector cytokines may be capable of thymic emigration. In particular, cells expressing high levels of CD24 have been suggested to dominate the recent thymic emigrant (RTE) pool of γδ-T cells^**12**,**13**^. In addition, a recent study utilizing the S1P1 signaling agonist FTY720 showed that γδ-thymocytes expressing high levels of CD24 and lacking expression of CD73 accumulated in the thymus following extended inhibitor treatment, again suggesting that many γδ-T cells may exit the thymus in a relatively immature state^**14**,**15**^. This suggests that the thymic environment may not be necessary for later stages of γδ-T cell maturation.

In this study, we performed an extensive analysis of γδ-thymocyte emigration from adult thymus. We find that γδ-RTEs in adult mice are largely immature CD73^-^ or immature CD73^+^ γδ-T cells that are capable of completing maturation in the periphery. A second stream of γδ-T cells completes maturation in the thymus before emigrating to the periphery. Furthermore, we identified a mature population of heterogeneous γδ-thymocytes that resided long term in the thymus and was generated during both the fetal/neonatal and adult periods of γδ-T cell development.

## Results

### Differential GFP expression in immature and mature γδ-thymocytes and T cells in adult Rag2pGFP mice

Within the thymus, there is a population of ‘mature’, CD24^-^, γδ-thymocytes that can produce cytokines such as IL-17 and IFNγ^**16**^. Considering that it is the most mature αβ-thymocytes that emigrate from the thymus^**17**^, one might predict that the mature, CD24^-^ population of γδ-thymocytes is exported from the thymus into the periphery as recent thymic emigrants (RTE). However, for other T cell lineages, such as iNK-T cells, it appears that immature precursors emigrate from the thymus^**18**^. In an attempt to directly examine emigration of γδ-thymocytes in adult mice, we utilized the Rag2pGFP model. Since GFP expression is under control of the Rag2 promoter in Rag2pGFP mice, intracellular GFP can be used as an inverse correlate of time passed since V(D)J recombination. We reasoned that we could track γδ-thymocyte development and emigration using these changes in GFP expression similar to previous studies examining αβ-T cell and iNK-T cell development^**17**,**18**^. We harvested the thymus and spleen from adult Rag2pGFP mice, gated γδ-TCR^+^CD3^+^ cells into sub-populations using CD24 and CD73, and measured GFP expression. Using DN3 thymocytes as a control, we found most immature CD73^-^ or immature CD73^+^ γδ-thymocytes expressed GFP, albeit at decreased levels when compared to DN3 thymocytes (Figure 1A,B). However, most mature γδ-thymocytes expressed drastically lower levels of GFP suggesting they are an aged product of immature CD73^-^ or CD73^+^ precursors. Of note, the GFP MFI among immature CD73^-^ or CD73^+^ γδ-thymocytes was similar, which suggests that they are similar in “age” (Supplementary 1A). In contrast, γδ-splenocytes showed a stepwise decrease in GFP expression accompanying maturation with most splenic mature γδ-T cells being GFP^-^ (Figure 1 A, B). No deviation from the general patterns correlating GFP expression and maturity in bulk γδ-thymocytes or splenocytes were found among the Vγ1.1 or Vγ2 subsets (Supplementary 1 A, B). In the spleen, the majority of GFP^+^ γδ-splenocytes were CD24^+^ and did not produce IFNγ or IL-17 following PMA/Ionomycin stimulation (Figure 1C, Supplemental 1C).

**Figure 1.**
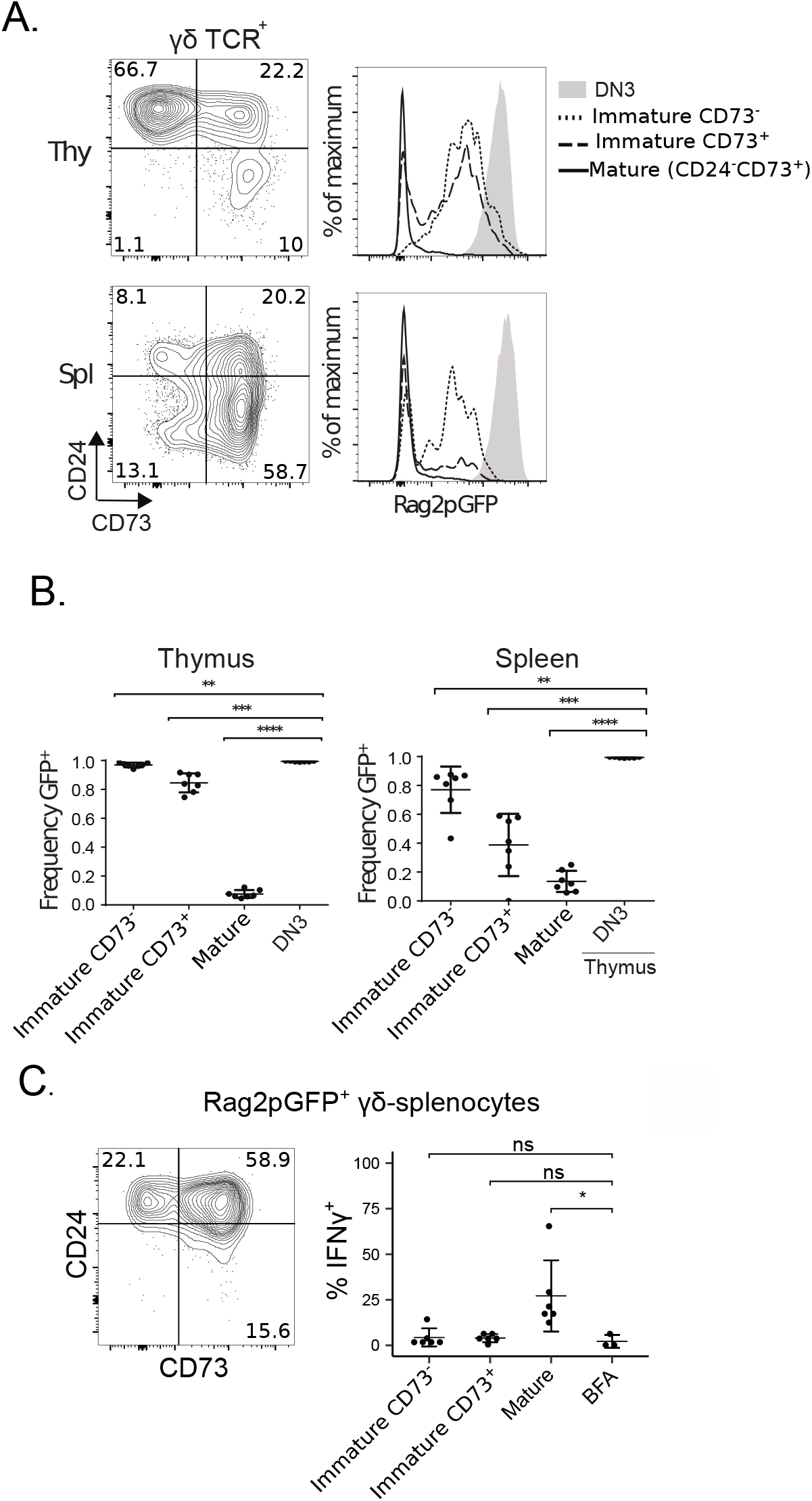
Differential GFP expression in immature and mature γδ-thymocytes and -splenocytes in adult Rag2pGFP mice. A. CD73 x CD24 gating of γδTCR+ thymocytes and splenocytes (left) from adult Rag2pGFP mice as immature CD73-(CD24+CD73-), immature CD73+ (CD24+CD73+), and mature (CD24-CD73+). GFP expression in gated sub-populations (right). B. Frequency of GFP+ cells in the subpopulations from A. C. CD73 x CD24 gating of GFP+ γδ-splenocytes (left), and IFNγ production of gated subpopulations after 4 hour PMA/Ionomycin stimulation (right). Mature splenocytes treated with BFA only used as control. Results shown are representative of at least 2 independent experiments. *p < 0.05, **p < 0.01, ***p < 0.001, ****p < 0.0001, ns = not significant.

Since GFP expression in the Rag2pGFP model ceases following successful V(D)J recombination, GFP detection may be influenced by the rate of cellular proliferation. Therefore, we used BrdU incorporation and Ki67 expression to examine γδ-thymocyte proliferation among immature CD73^-^, immature CD73^+^, and mature γδ-thymocytes. Using both BrdU and Ki67 as readouts of proliferation, mature γδ-thymocytes showed significantly lower frequencies of BrdU^+^ and Ki67^+^ cells compared to either immature CD73^-^ or CD73^+^ γδ-thymocytes (Supplementary 2A-B). Similar results were obtained when examining splenic γδ-T cells (Supplementary 2C-D).

Collectively, these data suggest that in adult mice, at least some of the peripheral γδ-T cell pool is composed of immature CD73^-^ or CD73^+^ cells and that these cells are likely RTEs. Additionally, low levels of GFP expression in mature γδ-thymocytes did not appear to be due to high levels of proliferation within this fraction, supporting the idea that these cells are not recently derived from immature thymic progenitors.

### Intra-thymic labeling reveals a heterogeneous pool of γδ-RTE, dominated by immature CD24^+^ γδ T cells

Similar to previous findings from Kelly et al.(1993)^**12**^, our data from the Rag2pGFP model suggested that CD24^+^ populations of γδ-thymocytes exit the thymus and traffic to secondary lymphoid organs. However, to determine the precise identity of the γδ-thymocyte RTE populations, we performed intra-thymic biotin labeling experiments in Rag2pGFP mice. Animals received intra-thymic biotin injections and biotin^+^ cells were magnetically enriched from spleens of recipient mice 24 hours after biotin administration. We used streptavidin-conjugated fluorophores to examine biotin^+^ (RTE) and biotin^-^ (non-RTE) γδTCR^+^CD3^+^ cells for expression of CD24 and CD73 to evaluate the composition of these fractions. Despite mature γδ-T cells making up the majority of the non-RTE splenic γδ-T cell population, the RTE fraction was largely composed of immature CD73^-^ or CD73^+^ γδ-T cells, with a minority of mature cells detected (Figure 2A-C). Furthermore, most γδ-T cell RTEs were GFP^+^ while the non-RTEs were GFP^-^. Since a similar CD24/CD73 pattern was found among biotin labeled γδ-splenocytes as in GFP^+^ γδ-splenocytes in Rag2pGFP mice (Figure 1C), we concluded that the majority of γδ-RTEs are in immature stages of development.

**Figure 2.**
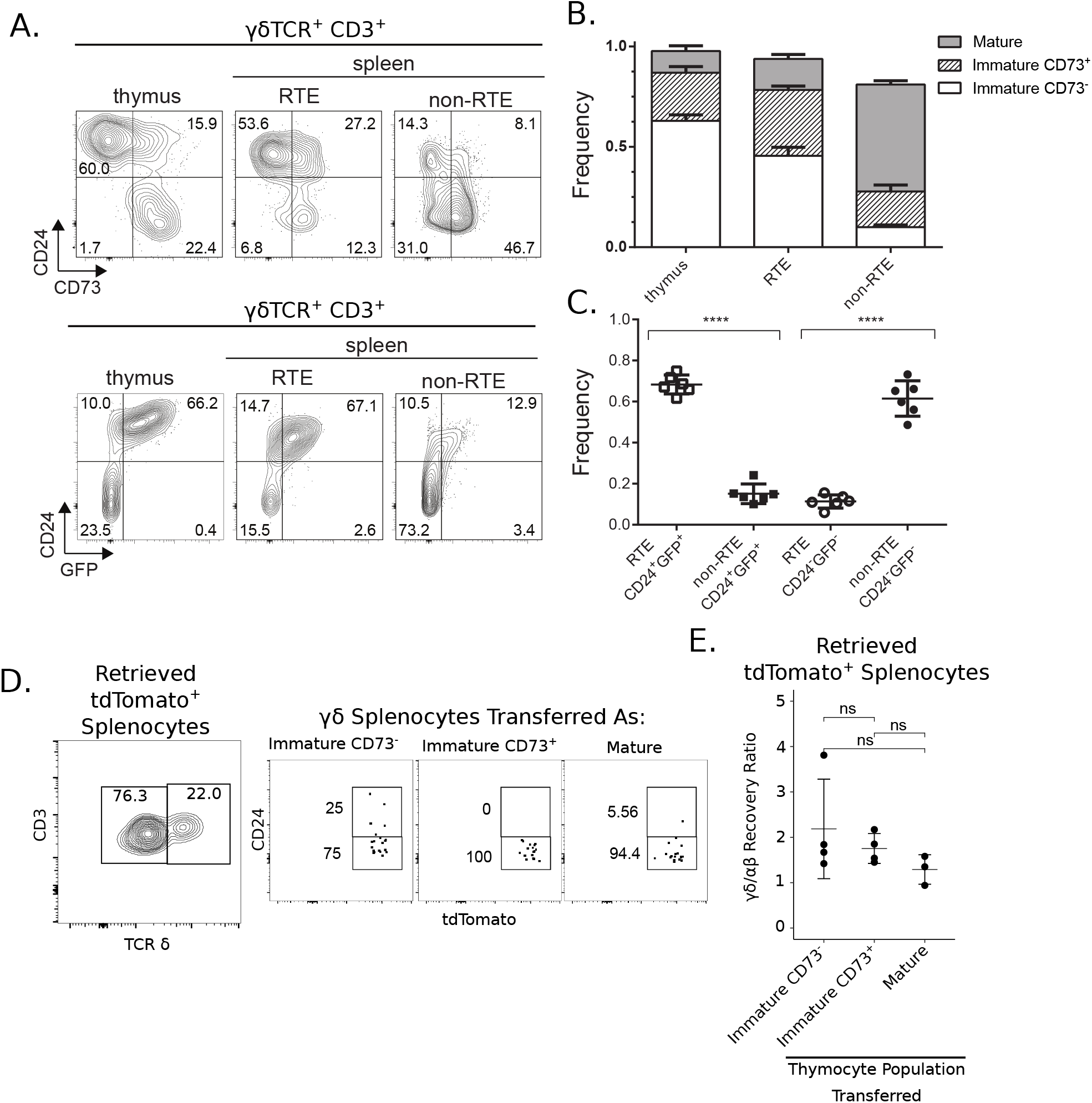
Most γδ-RTEs are immature and can mature in the periphery. A. Gating after enrichment of streptavidin-positive cells and subsequent analysis of CD73 x CD24 expression (top) and GFP x CD24 expression (bottom). B. Frequency of streptavidin-positive splenocytes by CD73 x CD24 subpopulations, in comparison with their unbound counterparts. C. Frequency of streptavidin-positive splenocytes by GFP x CD24 subpopulations, in comparison with their unbound counterparts. D. Gating of retrieved tdTomato+ γδ-splenocytes from adoptive transfer recipients (left) and representative plots of CD24 expression. E. Relative recovery of tdTomato+ γδ-T cells compared to tdTomato+ αβ-T cells from adoptive transfer recipients on day 3. Calculated as: (% of recovered tdTomato+ splenocytes that were γδ-T cells) / (% of adoptively transferred tdTomato+ cells that were γδ-thymocytes). Results shown are representative of at least 2 independent experiments. *p < 0.05, **p < 0.01, ***p < 0.001, ****p < 0.0001, ns = not significant.

### Some γδ thymocytes of all maturity levels are emigration competent

Expression of the egress receptor S1P1, and its transcriptional regulator KLF2, has previously been shown to be required for γδ-thymocyte trafficking out of the thymus, and we hypothesized that differences in thymic egress between immature CD73^-^, immature CD73^+^, and mature sub-populations may be influenced by expression of these molecules. Consistent with finding immature CD73^-^ or CD73^+^ and mature populations in the RTE pool, all three populations of γδTCR^+^CD3^+^ cells showed similar expression patterns of cell surface S1P1 (Supplementary 3A-C). Additionally, we sorted immature CD73^-^, immature CD73^+^, and mature γδ-thymocytes, and performed qPCR to measure transcript levels of *s1pr1* and *klf2*. Transcript levels of *s1pr1* and *klf2* were similar between the immature and mature γδ-thymocyte subpopulations (Supplementary 3D), suggesting that a similar fraction of all the different γδ-thymocytes subsets were competent for thymic export.

### γδ RTEs can mature in the periphery

We next sought to clarify the fate of immature γδ-RTEs. We hypothesized that either immature CD73^-^ or CD73^+^ γδ-RTEs continue maturation in the periphery, or they cannot survive in the periphery and only the minority of mature γδ-RTEs feed the peripheral γδ-T cell population. To test this, we generated mice where γδ-thymocytes could be inducibly labelled with a fluorescent reporter protein. Rosa26 lox-STOP-lox tdTomato mice were crossed with TCRδ-Cre^ERT2^ mice to allow for expression of tdTomato in γδ-T cells following tamoxifen administration. tdTomato labelled immature CD73^-^, immature CD73^+^, and mature γδ-thymocytes were purified by FACS and adoptively transferred into wild-type (WT) mice. Additionally, we co-adoptively transferred an equal number of mature tdTomato^+^ αβ-splenocytes as a reference population to compare survival/retrieval rates. After 3 days we harvested the spleen of recipient mice and identified the adoptively transferred populations by tdTomato expression. Most adoptively transferred immature CD73^-^ or CD73^+^ γδ-thymocytes were CD24^-^ at the time of retrieval suggesting continued maturation in the periphery (Figure 2 D). Analysis of the survival efficiency of transferred populations demonstrated that immature CD73^-^ or CD73^+^ and mature γδ-thymocytes survived for at least 3 days in the periphery at a rate as high or greater than transferred αβ-splenocytes (Figure 2 E). These findings suggest that immature CD73^-^ and CD73^+^ γδ-thymocytes can survive and mature in the periphery, and support the possibility that immature CD73^-^ or CD73^+^ γδ-RTEs contribute to the peripheral mature γδ-T cell pool.

### Parabiotic mice show minimal recirculation of γδ-T cells to the thymus, despite active circulation of γδ-splenocytes

Since mature γδ-T cells were a minority of γδ-RTEs and did not appear to be recently derived from immature γδ-thymocyte progenitors based upon GFP expression in the Rag2pGFP model, we next sought to investigate the dynamics of the mature γδ-thymocyte subset. One potential explanation for the relatively aged phenotype of mature γδ-thymocytes might be that this population is derived from recirculation of mature peripheral γδ-T cells. To address this possibility, we established parabiotic pairs of mice for 30 days where one mouse was CD45.1^+^ and its partner was CD45.2^+^. As expected, conventional CD4^+^ and CD8^+^ T cells from the spleen were nearly equally derived from each of the parabiotic partners (Figure 3A,B). Similar to conventional αβ-T cells, splenic γδ-T cells showed robust recirculation, with ∼60% of cells being host-derived on average (Figure 3A,B). In contrast to splenic T cell populations, thymocytes from both the αβ and γδ lineages showed very little contribution from the parabiotic partner. In particular, mature γδ-thymocytes showed significantly higher frequencies of host-derived cells compared to γδ-splenocytes (Figure 3B). This demonstrates that recirculation of mature peripheral γδ-T cells does not make a large contribution to the “aged” phenotype of mature γδ-thymocytes seen in Rag2pGFP mice.

**Figure 3.**
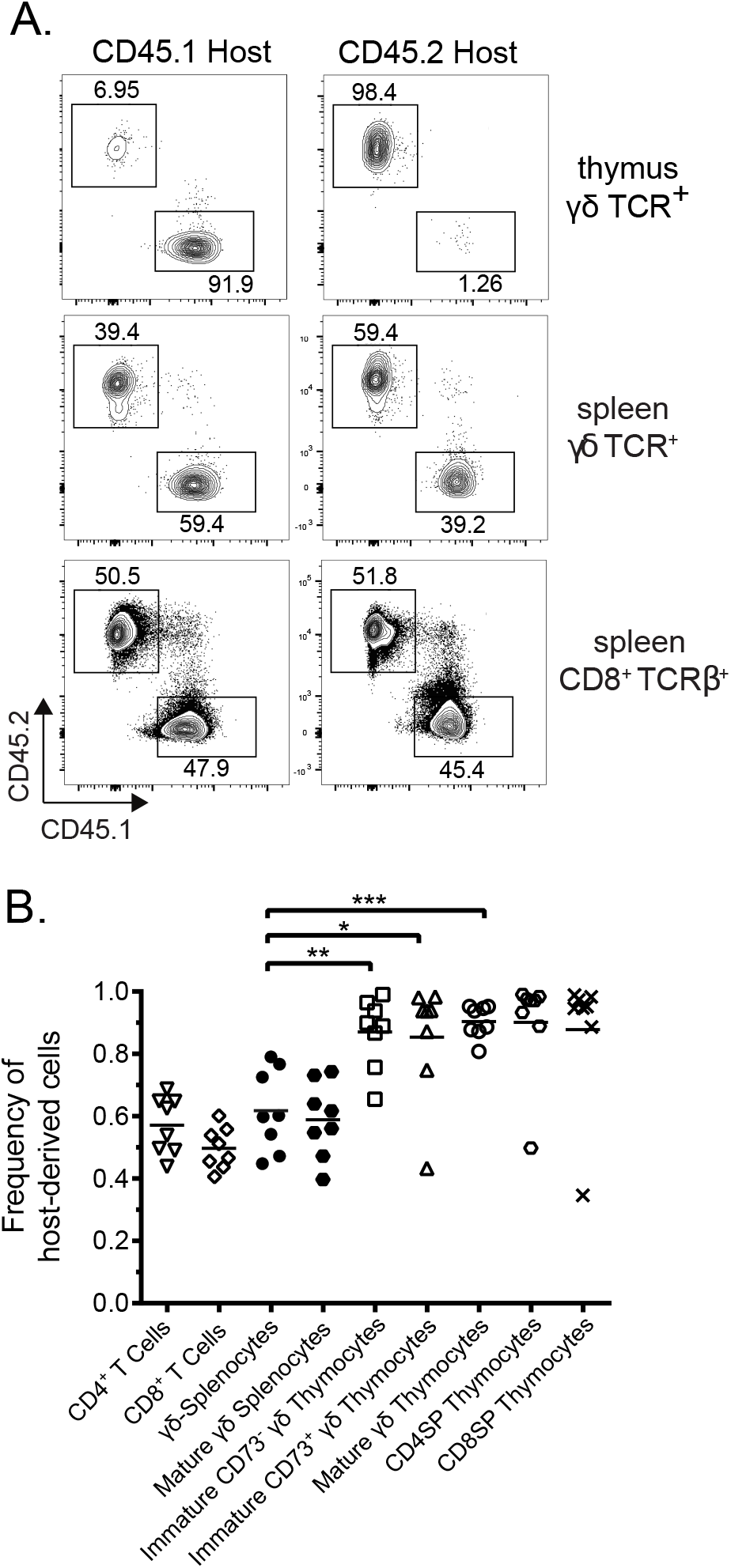
Peripheral γδ-T cells do not contribute to the γδ-thymocyte population. A. Flow cytometry profiles of CD45.1 vs CD45.2 derived cells in either host for both thymus and spleen after parabiosis. Splenic CD8+ T cells are presented as a reference. B. Frequency of host-derived cells among thymocyte and splenocyte populations. Results shown are representative of at least 2 independent experiments. *p < 0.05, **p < 0.01, ***p < 0.001, ns = not significant.

### A heterogeneous population of mature γδ thymocytes resides in the thymus long-term

To determine if the “aged” mature γδ-thymocyte population are longer-term resident cells, we utilized the TCRδ-Cre^ERT2^ Rosa26 lox-STOP-lox tdTomato mice. We reasoned that a pulse of tamoxifen followed by a chase of variable amounts of time would allow us to track the development and the emigration of γδ-thymocytes, and study any resident γδ-thymocytes that remained at the end of the chase. Tamoxifen was administered i.p. on days -1 and 0 and tdTomato expression in γδ-thymocytes was examined on days 2, 6, 15, 30, and 60. On day 2, tdTomato expression was observed in a fraction of all γδ-thymocytes and in all γδ-thymocyte subsets (Figure 4A). As expected, within the tdTomato^+^ γδ-thymocyte population, the frequency of both the immature CD73^-^ and CD73^+^ subsets decreased over time, while the frequency of the mature fraction increased over time (Figure 4A). Within the mature γδ-thymocyte population, the frequency and number of cells that expressed tdTomato was relatively constant for 60 days (Figure 4B). Additionally, the tdTomato^+^ mature γδ-thymocytes present on day 30 displayed a CD44^hi^ phenotype and both CD27^lo^ and CD27^hi^ subpopulations were present (Supplementary 4A). We consistently found a CD25^lo^ subpopulation of residents that were primarily CD27^-^ (Supplementary 4B). Mature day-30 tdTomato^+^ γδ-thymocytes showed substantial populations of Vγ1.1^+^, Vγ2^+^, and Vγ1.1^-^ Vγ2^-^ cells (Supplementary 4C). These findings demonstrate a heterogeneous population of longer-term resident mature γδ-thymocytes.

**Figure 4.**
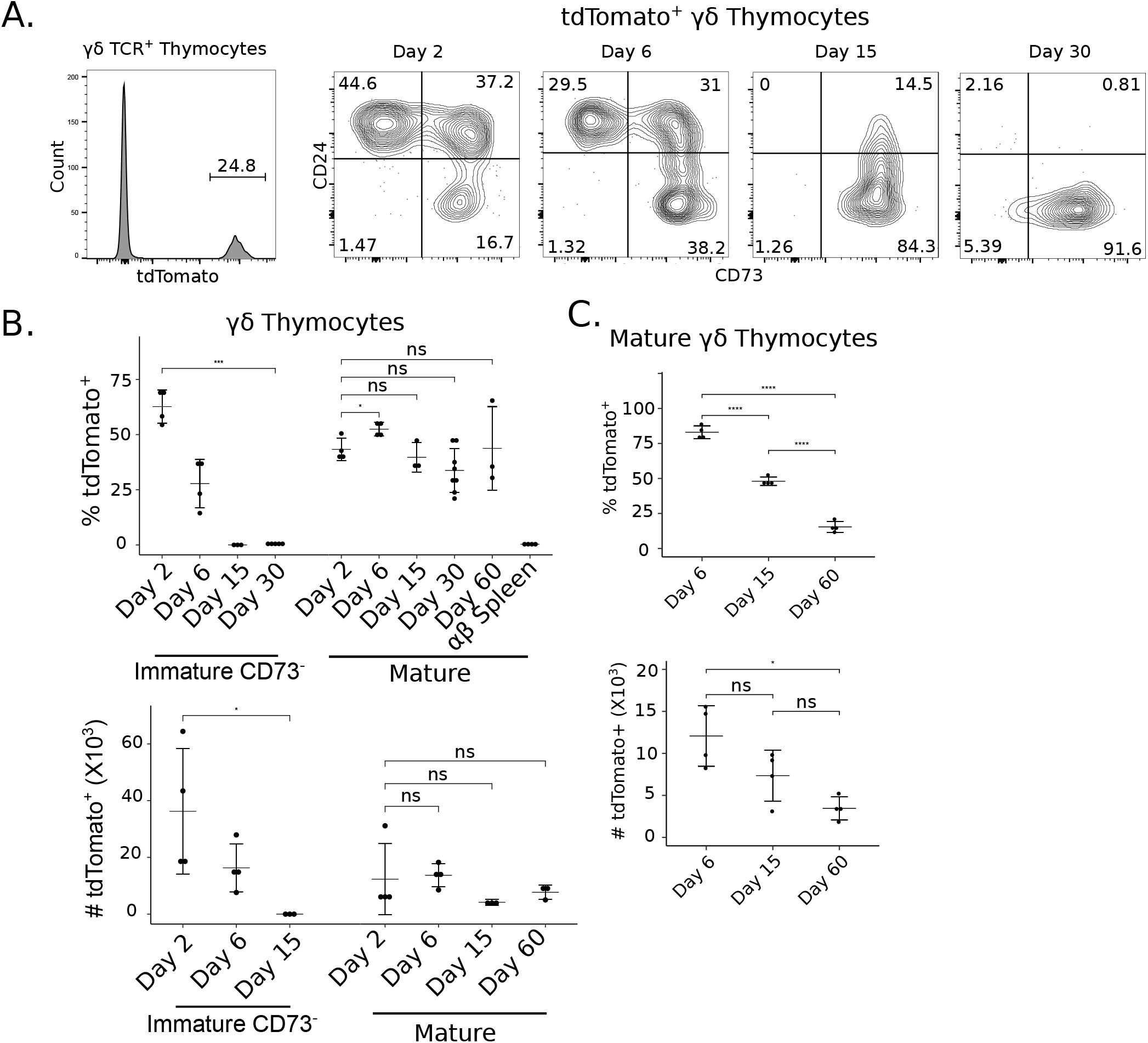
Fetal-derived mature γδ-thymocytes are thymic residents, whereas adult-derived γδ-thymocytes are lost from the thymus over time. A. Gating of tdTomato+ γδ-thymocytes (left), and CD73 x CD24 flow cytometry plot of tdTomato+ γδ-thymocytes over chase time (right). B. Frequency (top) and absolute number (bottom) of tdTomato+ immature CD73-and mature γδ-thymocytes over chase time. αβ-splenocytes sampled on Day 2 post-tamoxifen used as control. C. Frequency (top) and absolute number of tdTomato+ mature γδ-thymocytes over chase in TCRδ-CreErt2+tdTomato+ > WT Bone Marrow Chimeras. Results shown are representative of at least 2 independent experiments. *p < 0.05, **p < 0.01, ***p < 0.001, ****p < 0.0001, ns = not significant.

### Resident γδ-thymocytes are generated throughout life

Some populations of tissue-resident γδ-T cells are only generated during the fetal/neonatal period^19^, so we questioned whether these thymic residents were a product of fetal/neonatal development and/or adult development. To address this question, we generated bone marrow chimera (BMC) mice by transferring TCRδ-Cre^ERT2^ Rosa26 lox-STOP-lox tdTomato^+^ bone marrow into lethally irradiated WT adult mice, and after 6 weeks we treated the BMCs with tamoxifen. Similar to the intact, adult mice, the tdTomato^+^ immature CD73^-^ and CD73^+^ proportions decreased rapidly in the BMC mice. While the frequency and number of tdTomato^+^ mature γδ-thymocytes decreased over time, a population still remained after 60 days (Figure 4C).

We also analyzed the expression of Vγ chains present on the tdTomato labelled BMC thymocytes present on day 60 to determine if the composition of these residents differed from those in non-BMC mice. We consistently found that the day 60 residents generated in BMCs strongly favoured Vγ1.1 use (Supplementary 4D), while those long term residents found in intact adult thymi expressed more diverse Vγ chains (Supplementary 4C). Collectively, it appears that a population of fetal/neonatally-generated mature γδ-thymocytes forms a stable, long-term resident population while an adult derived mature γδ-thymocyte population resides in the thymus for a considerable amount, but is ultimately lost over time, possibly via emigration.

## Discussion

The finding that some γδ-thymocytes possess the capacity to produce cytokines shortly after stimulation has resulted in tremendous interest in understanding how γδ-thymocyte development and acquisition of effector function are coordinated. Intertwined in this area of investigation is the linkage between γδ-thymocyte development and emigration into the periphery. In this study, we found that the γδ-RTE population is largely composed of functionally immature cells (CD24^+^) rather than the most mature subset (CD24^-^) in adult mice. A second stream of developing γδ-thymocytes seems to complete maturation in the thymus before emigration to the periphery. Additionally, we determined that a population of mature γδ-thymocytes resides long-term in thymus. These findings raise questions about the factors that regulate thymic emigration and maturation of γδ-T cells, as well as the roles of resident γδ-thymocytes.

Our experiments using Rag2pGFP mice and intrathymic labelling demonstrated that the majority of γδ-RTEs emigrate while functionally immature. This finding is in agreement with previous work that suggested that most γδ-RTEs are CD24 ^+ **12**,**13**,**20**^. Previous studies indicated that like αβ-thymocytes, γδ-thymocytes utilize a KLF2-S1P1 emigration program^**10**^. Recent experiments that inhibited thymocyte emigration with an S1P1 receptor agonist, FTY720, found an accumulation of CD24^+^ γδ-thymocytes suggesting, albeit indirectly, that functionally immature CD24^+^ γδ-thymocytes are emigration competent. Similarly, through the use of genomics approaches, both the Lauritsen group and Sagar et al. identified populations of CD24^+^ γδ-thymocytes with *s1pr1* expression again supporting functionally immature γδ-thymocytes as being competent to emigrate^**15**,**21**^. Finally, while we observe expression of S1P1 on the surface of all γδ-thymocyte subsets it is interesting that only approximately half of all γδ-thymocytes in each subset appeared to express S1P1. That KLF2 was only expressed by approximately 40% of all γδ-thymocytes supports the idea that only a fraction of each γδ-thymocyte maturity-level is emigration competent^**10**^. Indeed, the maturity distribution of γδ-RTEs seems to approximate the maturity distribution of bulk γδ-thymocytes (Figure 2A). However, the factors that underlie the decision to express S1P1 or not at a given maturity level remain unclear.

With the finding that immature γδ-thymocytes constitute the major source of RTE, understanding the fate of these cells post-emigration was critical. Adoptively transferred CD24^+^ (immature CD73^-^ or CD73^+^) γδ-thymocytes were both capable of persisting in the periphery for 3 days with most cells having down-regulated CD24 in that time frame. In fact, when using mature αβ-T cells as a comparator population, it appeared that the adoptively transferred immature γδ-thymocytes expanded in the periphery. In support of these populations expanding post-transfer, CD24^+^ γδ-thymocytes or splenocytes are actively cycling based on BrdU incorporation and Ki-67 expression. In recent work by Sagar et al., a CD24^+^ population that expressed *s1pr1* and *ccr9* was found in the thymus and a similar population that expressed *s1pr1* and ccr9 was identified in the periphery^**21**^. While no direct precursor-product relationship was proven in this study, these findings further support the export of immature γδ-thymocytes followed by maturation in the periphery. Based on recent scRNA-seq data that revealed a substantial CD24^+^ CD44^-^ S1PR1^+^ population of γδ-thymocytes and splenocytes, we hypothesize that these immature γδ-RTEs are primarily the source of circulating naive-like γδ-T cells^**21**^.

While some mature γδ-thymocytes are able to quickly produce IL-17 or IFNγ, the developmental kinetics of these populations was unclear. Since IL-17 producing γδ-thymocytes are thought to be generated only during the fetal/neonatal period, but that they can be found in the adult thymus suggests they are a resident and self-renewing population^**5**^. In adult Rag2pGFP mice, the vast majority of CD24^-^ γδ-thymocytes were GFP^-^. This loss of GFP expression was not due to cell division as very few CD24^-^ γδ-thymocytes incorporated BrdU during a 2h pulse and most were Ki-67^-^. Parabiosis experiments demonstrated that the CD24^-^ γδ-thymocyte population were not derived from circulating peripheral pools. Pulse-chase labelling of γδ-T cells to track a single cohort of γδ-thymocytes demonstrated that this mature population remains resident in the thymus for at least 60 days. Taken together these findings indicate that many mature γδ-thymocytes may be residents.

The resident mature thymocyte population had a CD73^+^ and CD44^+^ phenotype, which suggests that this population is an effector-mature product of TCR interactions^**9**,**22**,**23**^. Since the population uses Vγ1.1, Vγ2, and other Vγ segments, it seems that it is a heterogeneous population that develops both during fetal/neonatal and adult life^**1**,**2**^. Indeed, the very presence of Vγ segments other than Vγ1.1. and Vγ2 suggests that some of these mature γδ-thymocytes were generated in the fetal/neonatal period of life. In addition, previous experiments with thymic grafts have shown that donor derived, and likely fetal-derived, γδNKTs reside in the thymus long-term^**24**^. Extending the pulse-chase labelling approach to study γδ-thymocyte development from strictly adult progenitors, our BMC experiments demonstrated that adult-derived mature γδ-thymocytes spend a considerable amount of time in the thymus, but ultimately decay over time with a half-life of approximately 10 days (Figure 4C). Therefore, we hypothesize that a minority of mature γδ-thymocytes are generated in adulthood from developing γδ-thymocytes and reside in the thymus for a relatively long period of time before emigrating to the periphery.

What function(s) do resident γδ-thymocytes perform *in situ*? During embryonic development of the thymus, Vγ3^+^ DETC progenitors provide RANKL signals to induce the initial generation of AIRE^+^ mTEC^**25**^. Whether resident γδ-thymocytes in adult mice similarly provide factors that ensure proper thymic homeostasis is unclear. Since the thymus is largely an epithelial tissue, it seems possible that, as in other epithelial tissues like skin, thymic resident γδ-thymocytes capable of producing these cytokines could act as sentries against thymic infection. Considering the risk of central tolerance to pathogens that thymic infection could bring^**26**^, it would make sense for resident γδ-thymocytes to evolve to coordinate a rapid immune response to thymic infection. In addition, mounting evidence suggests that tissue-resident γδ-T cells play substantial regulatory roles in epithelial tissues, such as tissue repair via amphiregulin production in gingiva^**27**^. In fact, recent experiments demonstrated elevated amphiregulin expression in presumably resident Vγ4^+^ γδ-thymocytes without overt activation or constitutive IL-17 production^**28**^. We hypothesize that resident γδ-thymocytes may perform regulatory roles specific to thymus epithelial homeostasis. Indeed, some evidence suggests that the CD25^lo^ population of γδ-thymocyte residents is fetal-derived and expands in response to thymic stress^**29**^.

In summary, our findings suggest a model of γδ-T cell developmental dynamics wherein there are two streams of developing γδ-thymocytes in adult thymus. Most γδ thymocytes emigrate while immature and CD73^-^ or CD73^+^, then complete maturation in the periphery. A second population remains to complete maturation in the thymus before emigration to the periphery. In addition, a third population of fetal-derived γδ-thymocytes are permanent thymic residents.

## Materials And Methods

### Mice

All mice were bred and maintained in our colony at the University of Alberta and maintained on the C57BL/6 background. Rag2pGFP^**30**^ mice were generously provided by Dr. Pamela Fink. *Tcrd*^*tm1*.*1(cre/ERT2)Zhu*^/J (TCRδ-Cre^ERT2^) mice were obtained from The Jackson Laboratory and *Gt(ROSA)26Sor*^*tm14(CAG-tdTomato)Hze*^/J (conditional tdTomato reporter mice) were provided by Dr. Jeffrey Tessem (BYU). TCRδ-Cre^ERT2^ tdTomato reporter mice were generated by intercrossing. OT-I CD4Cre and OT-I CD8Cre tdTomato mice were similarly generated. Mice were analyzed between 8-14 weeks of age. All mice were treated in accordance with protocols approved by the University of Alberta Animal Care and Use Committee and the Institutional Animal Care and Use Committee at the University of Minnesota.

### Bone Marrow Chimeras

Donor TCRδ-Cre^ERT2^ mice were depleted of T cells by injection of 100ug of purified 30H12 (anti-Thy1.2) antibody (BioXcell) on days -2 and -1. Recipient wildtype mice were lethally irradiated with 2X 450 Rad separated by 1-4 h on day -1 and transplanted on day 0 with 5-10×10^6^ donor bone marrow cells. Mice were provided novotrimel in their drinking water for 28 days post-transplant.

Chimeric mice administered tamoxifen 8 weeks post-transplant.

### Antibodies and Flow Cytometry

Fluorochrome-conjugated and biotinylated antibodies were purchased from BD Biosciences, BioLegend, and eBioscience. Rat anti-mouse S1PR1 primary antibody was purchased from R&D Systems, biotinylated anti-rat secondary antibody from eBioscience (#13-4813-85), and BV-711-Streptavidin-conjugated tertiary from BD Biosciences (#563262). Live/Dead staining was done using Live/Dead Fixable Aqua Dead Cell stain kit (Thermo Fisher Scientific: L34957). Permeabilization for intracellular antigen staining was done using either BD Cytofix/Cytoperm (BD Biosciences) or the Foxp3 Staining Buffer Set (eBioscience). Cells were stained by treatment with antibody cocktails in FACS buffer (PBS, 1% FCS, 0.02% sodium azide, 1mM EDTA) for 30 minutes on ice in the dark. Cells were washed twice between staining steps. S1P1 staining was done in S1P1 Staining Buffer (PBS, 1% charcoal-stripped FBS (Thermo Fisher Scientific: A3382101), 1 mM EDTA), and stained for 90 minutes on ice. Blocking steps were done with Normal Mouse Serum (Cedarlane: CLS303) and/or Normal Rat Serum (STEMCELL Technologies: 13551). Flow cytometry was performed on LSR Fortessa (BD Biosciences) or Attune NXT (Invitrogen), and data was analyzed with FlowJo V10.6.2 (BD Biosciences).

### Cytokine Induction

Bulk thymocytes or splenocytes were treated with Brefeldin A (Invitrogen: 00-4506-51), PMA 20ng/mL, and ionomycin 1 μg/mL incubated at 37° C 5% CO_2_ for 4 hours in RP10 media. Cells were harvested and processed for flow cytometry as described above.

### Intrathymic Injection

Intrathymic injection of biotin was performed as described by Blair-Handon et al^31^, Briefly, Rag2pGFP mice were anesthetized with isoflurane, hair in the thoracic region was removed and the thymus was visualized with the VEVO3100 imaging system (Visualsonics). Each thymic lobe was injected with 10uL of a 1mg/mL solution of sulfo-NHS-biotin in PBS (Pierce). 24 h post-injection the thymus and spleen were removed. Biotinylated splenocytes were enriched using the EasySep Biotin Positive Selection Kit (STEMCELL) and thymocytes and splenocytes were processed for flow cytometric analysis as described above.

### Tamoxifen Treatment

Mice between the ages of 8-12 weeks were treated with tamoxifen (Sigma-Aldrich: T5648-16) dissolved in corn oil to 20 mg/mL. Doses of 100 microliters were given on days -1 and 0 I.P., then lymphoid organs were analyzed on days 2, 6, 15, 30, or 60 for the presence of tdTomato^+^ γδ T cells.

### Adoptive Transfers

On day 2 post-tamoxifen treatment, induced tdTomato expressing thymocytes were labelled with biotinylated anti-CD4 and -CD8 antibodies and were enriched by immunoseparation of bulk thymocytes with Easysep Streptavidin Rapidspheres (Stem Cell Technologies) to reduce CD4^+^ and CD8^+^ cells. FACS was used to sort tdTomato+ cells that were CD24^+^/CD73^-^, CD24^+^/CD73^+^, or CD24^-^/CD73^+^. Sorted populations were transferred IV into wild-type mice along with an equal number of CD4Cre^+^ tdTomato^+^ competitors. Mice were analyzed 3 days post-transfer.

### BrdU

Thymocyte and splenocyte BrdU incorporation was assessed by flow cytometry. Mice were injected with 1mg of BrdU (Sigma) in PBS i.p. and euthanized 2h later. Thymus and spleen were harvested and single cell suspensions were incubated with antibody cocktails against cell surface proteins as above. Cells were fixed and permeabilized with the BD Fix/Perm kit and treated with 0.3mg/mL DNase I for 1h at 37°C. Incorporated BrdU was detected using anti-BrdU APC (Phoenix Labs).

### Parabiosis

Parabiosis surgeries were performed as previously described with some small alterations^**32**^. Briefly, CD45.1 and CD45.2 mice were anesthetized with ketamine, flank hair was shaved and the skin was wiped with ethanol. A lateral incision was made on the abdomen on the right side of one mouse, and a corresponding incision was made on the left side of the partner mouse. Surgical clips were used to join the mice at the incision and the skin surface was disinfected with betadine. Sustained release buprenorphine and bupivicaine were given to the mice after anesthesia, but before surgery. Parabiotic pairs were analyzed 30 days after surgery.

### RT-PCR

CD3^+^TCRγδ^+^ Immature (CD24^+^CD73^-^), Committed (CD24^+^CD73^+^) or Mature (CD24^lo^CD73^+^) thymocytes and CD4^+^CD8^+^ DP thymocytes were purified by FACS (BD FACS Aria III) and the RNA was isolated using RNeasy micro kit (Qiagen) with on column DNase digestion. cDNA was synthesized using Superscript III reverse transcriptase kit (Invitrogen). qRT-PCR analysis was performed with PerfeCTa SYBR Green Fastmix (Quantabio) using the following primers from Sigma-Aldrich: *S1pr1* (5’-AGCTTTTCCTTGGCTGGAGAG, 5’-GTGTAGACCCAGAGTCCTGCG), *Klf2* (5’-CGTGTTGGACTTCATCCTCTC, 5’-CGGCTCCGGGTAGTAGAAG)

### Statistics

Statistical analysis was performed using R. Mean and Standard Deviation were calculated with the plyr package (v1.8.6). Error bars represent one standard deviation. Dot plots were generated and two-tailed unpaired student’s t tests were performed using the ggpubr package (v0.4.0). Asterisks represent statistically significant differences between groups annotated as P < 0.05 (*), P < 0.01 (**), P < 0.001 (***), P < 0.0001 (****).

## Acknowledgements

The authors thank Drs. Gabrielle Siegers and Heather Melichar for critical review of the manuscript. We are grateful to Dr. Jason Dyck and Jody Levasseur and the Cardiovascular Research Centre for assistance with the image-guided intrathymic injection procedure. Flow cytometry experiments were performed at The University of Alberta Faculty of Medicine & Dentistry Flow Cytometry Facility, which receives financial support from the Faculty of Medicine & Dentistry and Canada Foundation for Innovation (CFI) awards to contributing investigators. Health Sciences Laboratory Animal Services and Mr. Bing Zhang provided animal husbandry and technical assistance.

## Author contributions

K.J., D.P.G., H.W., S.C.J. and T.A.B. designed the experiments. K.J, D.P.G., H.W., L.H.C. and T.A.B analyzed the data. K.J., D.P.G. and T.A.B. wrote the manuscript. K.J., D.P.G., H.W., L.M.H.C., R.G.K., J.F.M. and R.C. performed the experiments.

