## Supplementary Figures for "γδ-Thymocyte Maturation and Emigration"

### Supplementary Figure 1

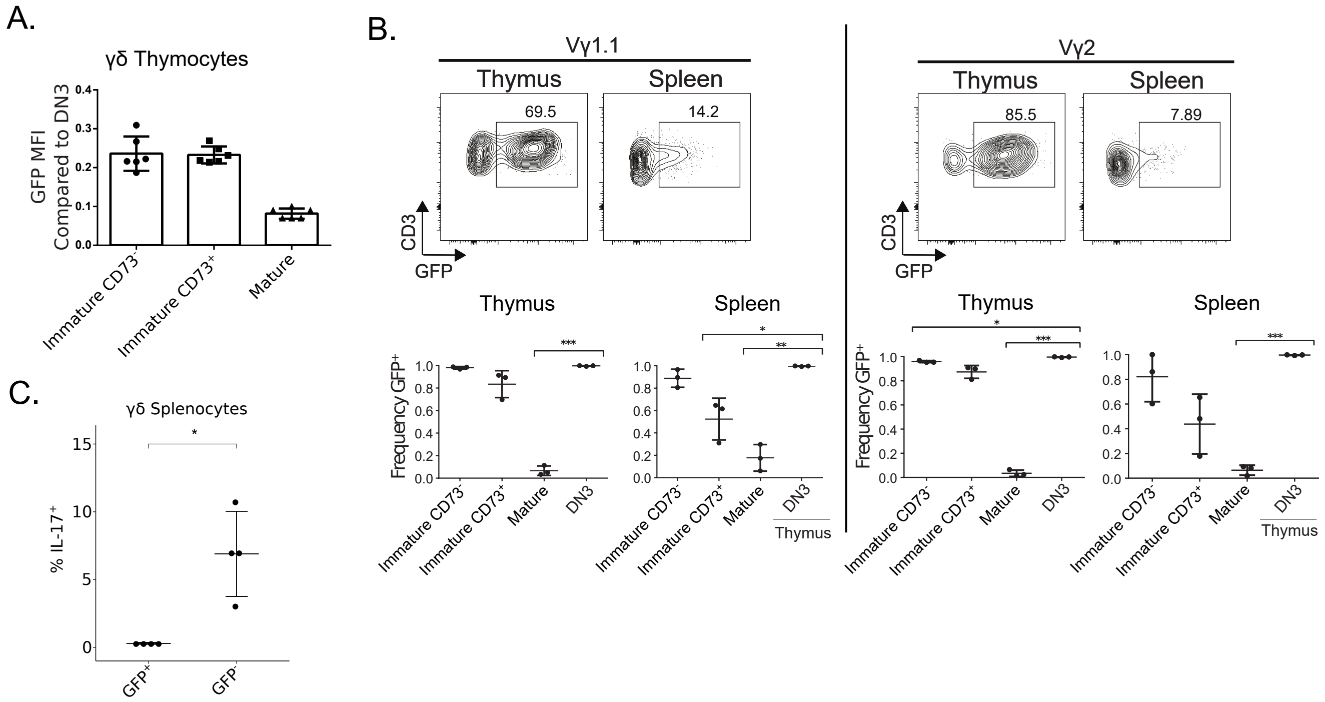


Supplementary 1. Most immature γδ-thymocytes and -splenocytes are GFP+ and do not produce cytokines. A. MFI of GFP among γδ-thymocyte sub-populations. B. GFP positive gating of Vγ1.1+ and Vγ2+ cells from the thymus and spleen (top) and frequency of GFP expression by subpopulation in the thymus and the spleen (bottom). C. IL-17 production by GFP+ or GFP- γδ-splenocytes stimulated for 4 hours with PMA/Ionomycin. Results shown are representative of at least 2 independent experiments. *p < 0.05, **p < 0.01, ***p < 0.001, ****p < 0.0001, ns = not significant.

Supplementary Figure 2

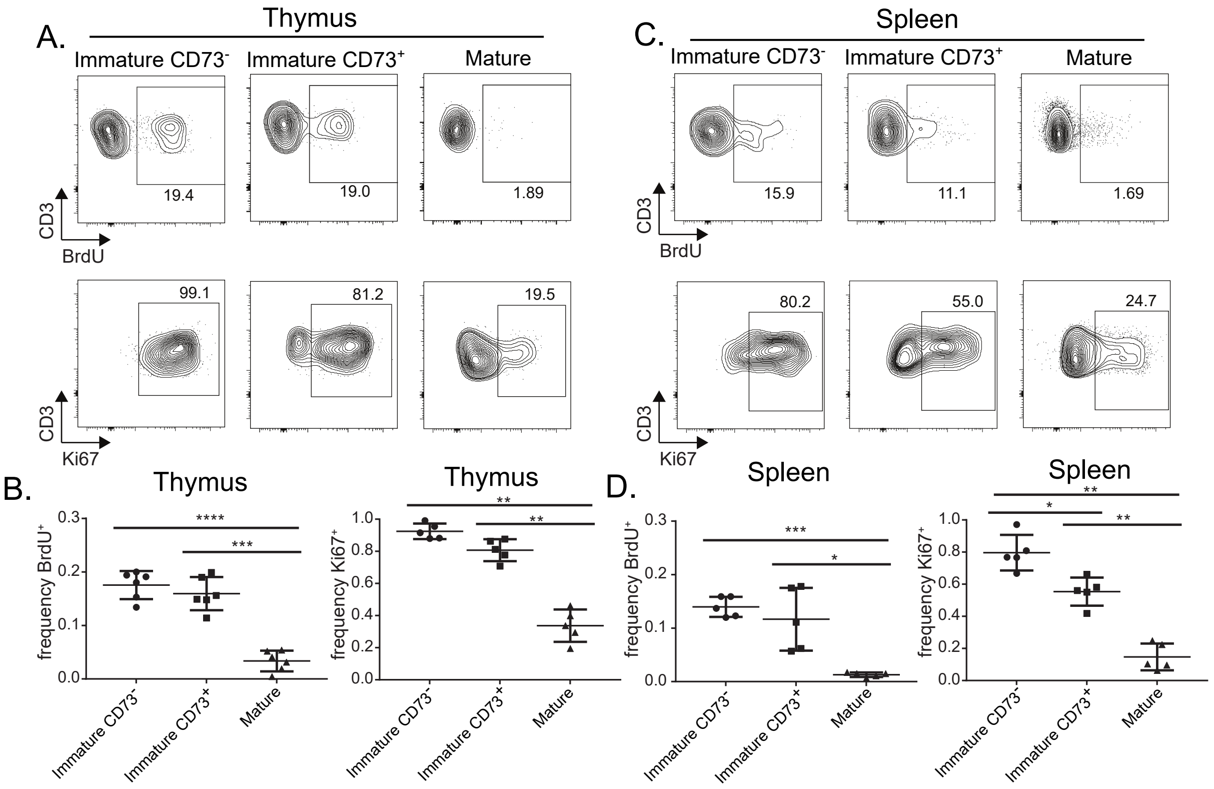


Supplementary 2. Proliferation of γδ-thymocyte and -splenocyte subpopulations. A. Representative flow cytometry profiles of BrdU incorporation (top) and Ki67 expression (bottom) in the indicated γδ-thymocyte subpopulations in thymus. B. Compilation of the frequency of BrdU+ (top) and Ki67+ (bottom) γδ-thymocyte subpopulations. C. Representative flow cytometry profiles of BrdU incorporation (top) and Ki67 expression (bottom) in the indicated splenic γδ-T cell subpopulations. D. Compilation of the frequency of BrdU+ (top) and Ki67+ (bottom) splenic γδ-T cell subpopulations. Results shown are representative of at least 2 independent experiments. *$p < 0.05$, **$p < 0.01$, ***$p < 0.001$, ****$p < 0.0001$, ns = not significant.

Supplementary Figure 3

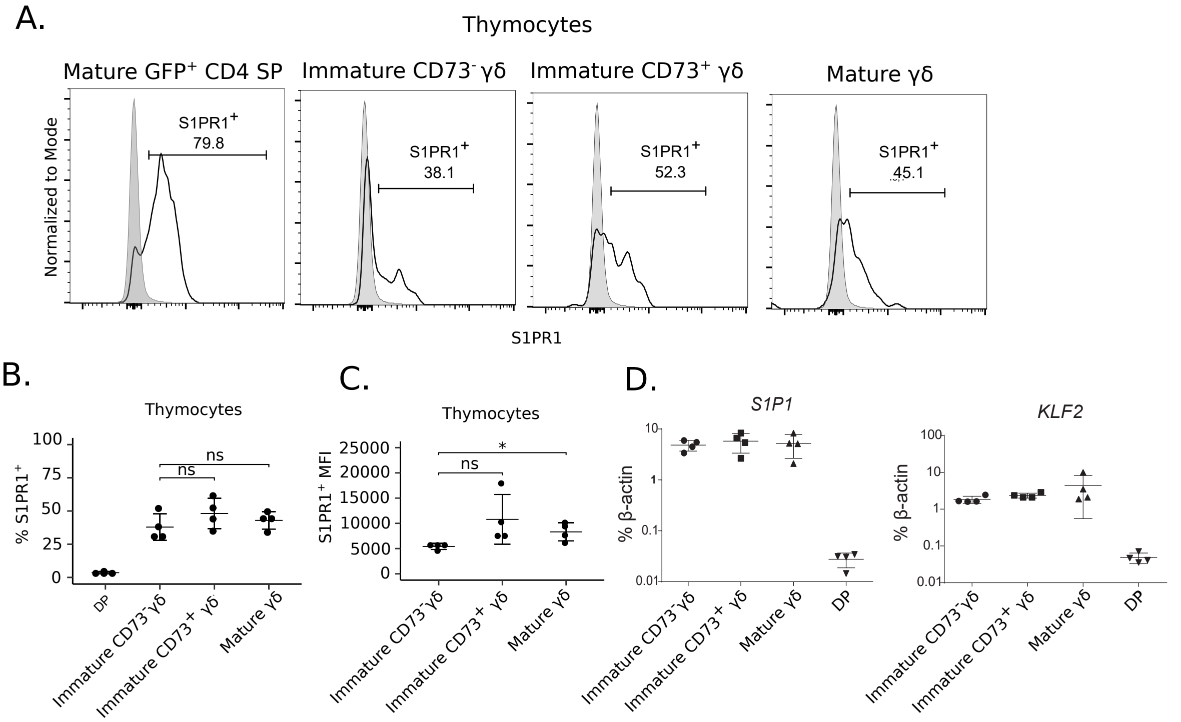


Supplementary 3. Expression of S1PR1 in γδ-thymocyte sub-populations. A. Histograms of S1PR1 expression among thymocyte subsets. The DP thymocyte negative control is shaded, and CD4SP thymocytes are depicted as positive control. B. Expression of S1PR1 among γδ-thymocyte subsets as measured by flow cytometry. CD4/CD8 DP population included as negative control. C. MFI of S1PR1+ γδ thymocyte subsets. D. RT-PCR for S1P1 and KLF2 expression among γδ-thymocyte sub-populations. Results shown are representative of at least 2 independent experiments. *$p < 0.05$, **$p < 0.01$, ***$p < 0.001$, ****$p < 0.0001$, ns = not significant.

Supplementary Figure 4

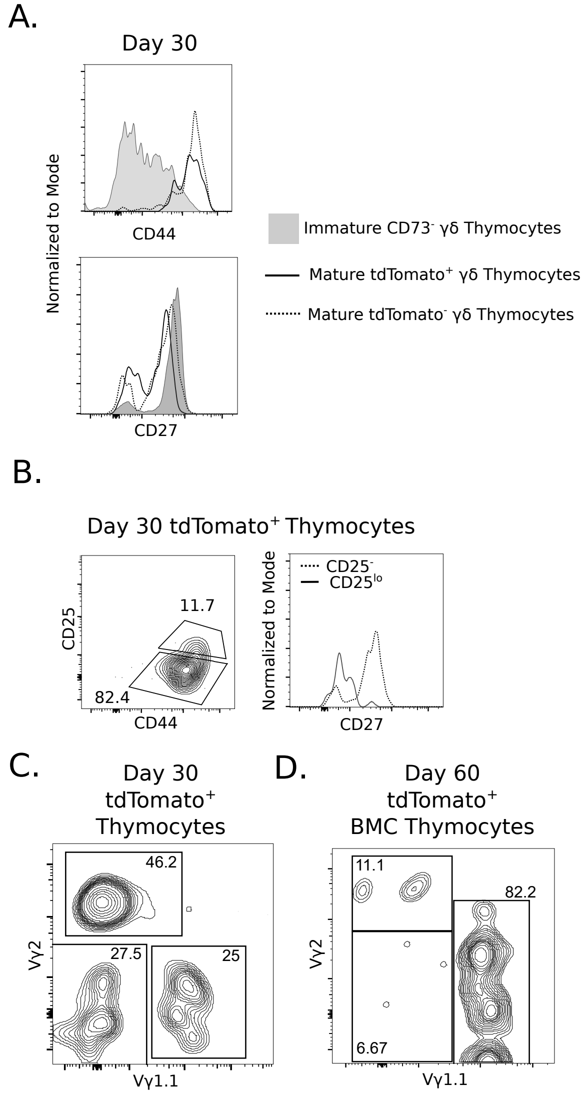


Supplementary Figure 4. Resident γδ-thymocytes are heterogeneous. A. Expression of CD44 and CD27 in the indicated population. B. Representative CD25 vs. CD44 expression in resident γδ-thymocytes (left) and CD27 expression on CD25- and CD25lo resident γδ-thymocyte subsets (right). C. Vγ usage within resident γδ-thymocytes at day 30 post-tamoxifen. D.Vγ usage within tdTomato labelled γδ-thymocytes at day 60 post-tamoxifen in TCRδ-CreERT2+ tdTomato+ -> WT BMCs. Results shown are representative of at least 2 independent experiments. *p < 0.05, **p < 0.01, ***p < 0.001, ****p < 0.0001, ns = not significant.
